# Decreased viability and neurite length in neural cells treated with chitosan-dextran sulfate nanocomplexes

**DOI:** 10.1101/616144

**Authors:** Laura N. Zamproni, Daniela Teixeira, Amanda Arnaut Alliegro, Ieda Longo Maugéri, Anne des Rieux, Marimelia A. Porcionatto

**Affiliations:** Neurobiology Lab, Department of Biochemistry, Escola Paulista de Medicina, Universidade Federal de São Paulo, Rua Pedro de Toledo 669, 3° andar, Vila Clementino, São Paulo, CEP: 04039-032; Division of Immunology, Department of Microbiology, Immunology and Parasitology, Escola Paulista de Medicina, Universidade Federal de São Paulo, Rua Botucatu, 862 quarto andar Vila Clementino, São Paulo, CEP: 04023-900; Université Catholique de Louvain, Louvain Drug Research Institute, Advanced Drug Delivery and Biomaterials, Avenue E. Mounier 73, 1200 Bruxelles, Belgium

**Keywords:** Ischemic stroke, CXCL12, Chitosan, Dextran sulfate, nanocomplexes

## Abstract

CXCL12 is a chemokine known to regulate migration, proliferation, and differentiation of neural stem cells (NSCs) and to play a neuroprotective role in ischemic stroke. Chitosan-dextran sulfate nanocomplexes (Ch/DS NC) are known nanoparticulated systems used to efficiently deliver heparin-binding factors. Here we evaluate Ch/DS NC as carriers for CXCL12 in a mouse model of stroke. Free CXCL12 reduced the size of the ischemic brain lesion. However, when Ch/DS NC were administrated, the stroke volume increased. Neurotoxic screening revealed that Ch/DS NC reduced neuronal viability, decreased the extension of neurites and impaired NSC migration *in vitro*. To the best of our knowledge, neurotoxicity of Ch/DS NC has not been reported and further screenings will be needed in order to evaluate the biological safety of these nanocomposites. Our results add new data on nanoparticle neurotoxicity and may help us to better understand the complex interactions of the nanostructures with biological components.

## Introduction

C-X-C chemokine ligand 12 (CXCL12), also known as stromal-derived factor-1 (SDF-1), is a chemokine that regulates migration, proliferation, and differentiation of neural stem cells (NSCs) within the developing central nervous system (CNS) (Li, Tang et al. 2015). Although there is some controversy (Pan, Yan et al. 2016, Cheng, Lian et al. 2017, Wu, Yu et al. 2017), there is increasing evidence showing that CXCL12 overexpression or delivery improves neurobehavioral recovery during post-ischemic stroke (Shyu, Lin et al. 2008, Li, Tang et al. 2015). Previous treatment of stroke with CXCL12 was shown to upregulate CXCL12/CXCR4 axis and enhance neurogenesis, angiogenesis, and neurological outcome (Shyu, Lin et al. 2008, Yoo, Seo et al. 2012, Kim, Seo et al. 2015). However, CXCL12 is prone to inactivation, denaturation, and degradation and improving its *in vitro* stability and *in vivo* half-life is essential for clinical applications (Zaman, Wang et al. 2016).

The CXCL12 protein structure has a heparin-binding site, which allows it to bind to heparan sulfate, form dimers that are less susceptible to protease inactivation, and interact with target cells via cell surface receptors. Since dextran sulfate (DS) has similar structural properties as heparin and heparan sulfate, it can bind CXCL12 in a way similar to that of its natural polymeric ligands. DS is a polyanion that can form polyelectrolyte complexes through electrostatic interactions with the polycation Chitosan (Ch) (Bader, Li et al. 2015).

Chitosan-dextran sulfate nanocomplexes (Ch/DS NC) have advantages for protein delivery compared to other nanoparticle systems, such as high hydrophilicity. Complexes are produced in water via charge-charge interactions, which pose minimal risk for denaturation or inactivation of the incorporated proteins. Additionally, binding to DS or Ch in the matrix of nanocomplexes provides dynamic immobilization and protection of the incorporated proteins (Bader, Li et al. 2015, Zaman, Wang et al. 2016).

In fact, CXCL12 associated with Ch/DS NC exhibit enhanced stability and sustained *in vivo* effects/retention (Zaman, Wang et al. 2016). When aerosolized into the lungs of rats, CXCL12 Ch/DS NC exhibited longer retention time than that of free CXCL12 (t_1/2_ = 20 h, and 3.2 h, respectively; 29% remaining after 72 h and 2% after 16 h, respectively). The aerosolized CXCL12 Ch/DS NC, but not free CXCL12, reduced pulmonary artery pressure in rats (Yin, Bader et al. 2013).

As far as we know CXCL12 Ch/DS NC have not been used in brain injuries and Ch/DS NC neurotoxicity has not been tested. In this work we evaluated Ch/DS NC as a carrier for CXCL12 in a stroke mouse model. However, we found evidence that Ch/DS NC increased stroke area, worsened mice motor deficits after ischemia, and decreased cell viability and neurite length *in vitro*.

## Materials and Methods

### Cells

Jurkat cells (human acute T lymphocytic leukemia cell line) were cultured in RPMI1640 medium (Cultilab,Campinas, Brazil) with 10% fetal bovine serum (FBS, Cultilab), 1% glutamine (Gibco, Waltham, USA), and 1% penicillin/streptomycin antibiotics (Gibco). N2a cells (murine neuroblastoma cell line) were cultured in high glucose DMEM medium (Gibco) with 10% FBS, 1% glutamine and 1% penicillin / streptomycin.

Adult neural stem cells (NSCs) were obtained from the subventricular zone (SVZ) of 45 days old C57BL/6 female mice according to previous described protocol (Adelita, Stilhano et al. 2017). Cells were cultured in DMEM/F 12 1:1 (v/v) (Gibco), supplemented with 2% B27 (Thermo Fischer, Waltham, USA), 20 ng/mL EGF (Sigma, St Louis, USA), 20 ng/mL FGF2 (Sigma), 1% penicillin/streptomycin (Gibco), and 5 μg/mL heparin. Cells were plated on a poly 2-hydroxyethyl metracrylate (poly HEMA) (Sigma) pre-coated 75 cm^2^ flask to avoid adhesion. Neurospheres formation takes up to 14 days to occur, and during this time culture medium was partially changed every 3 days.

All cells were kept at 37°C and 5% CO_2_.

### Nanocomplexes (NC)

#### Production of CXCL12 Ch/DS NC

CXCL12 Ch/DS NC were produced following a protocol described (des Rieux, Ucakar et al. 2011). Five µl of CXCL12 (1 µg/µL) (Peprotech, Rocky Hill, USA) were added to 50 µL of 1% DS (500 kDa, Fluka, Buchs, Switzerland) and stirred for 30 min. One hundred µL of 0.1% chitosan solution (Polysciences, Valley Road, Warrington) prepared in 1% acetic acid (pH 5) were added dropwise to the CXCL12-DS solution and stirred for 5 min. Five µL of 1 M zinc sulfate (Sigma) were added to the CXCL12-DS-Ch solution and stirred for 30 min. NC were centrifuged in an Eppendorf tube containing 10 µL of glycerol for 30 min at 14000 g. Supernatant was collected and frozen for further encapsulation efficiency and drug loading analysis. NC were suspended in 200 µL PBS and used immediately after production. Ch/DS NC were produced by substituting CXCL12 by 5µL of sterile water.

#### NC characterization

NC size was measured by NTA with a NanoSight (NanoSight, Salisbury, United Kingdom) and zeta potential were measured with Zeta sizer (Malvern Instruments, Malvern, United Kingdom). The shape and surface of the dried NC were observed by Scanning electron microscopy (SEM), using a Quanta 650 FEG microscope (FEI, Thermo Fischer, Waltham, USA).

#### Encapsulation efficiency (EE) and Drug loading (DL)

CXCL12 concentration in supernatants of CXCL12 Ch/DS NC was measured using Mouse CXCL12 DuoSet ELISA as per supplier instructions (R&D systems, Minneapolis, USA). Encapsulation efficiency and drug loading were calculated as follow:

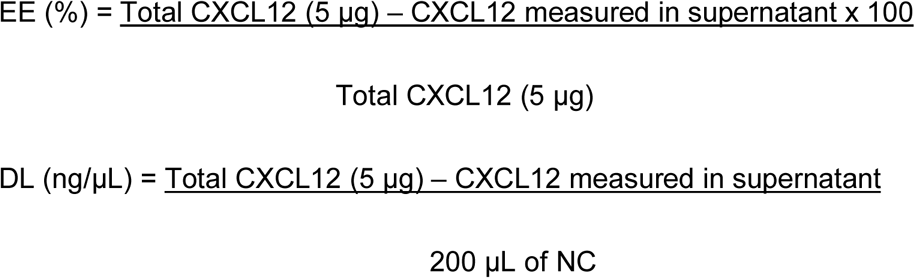

#### CXCL12 release

Five µL NC were suspended in 200 µl of PBS (n=3). Samples were incubated at 37 °C and at given times, were centrifuged at 14000 g for 30 min at room temperature. Supernatants were collected and frozen until further analysis. CXCL12 was quantified from the supernatants using Mouse CXCL12 DuoSet ELISA (R&D Systems).

#### Western Blot

Purified CXCL12 and CXCL12 Ch/DS NC were resolved by 15% sodium dodecyl sulfate (SDS)-polyacrylamide gel electrophoresis (Bio-Rad, Hercules, USA) and electro transferred to a polyvinylidene difluoride (PVDF) membrane (Merck Millipore, Burlington, EUA). After blocking with 5% bovine serum albumin (BSA; Sigma), the PVDF membrane was sequentially probed with specific primary antibodies (1:1000, anti-CXCL12, rabbit Ig G, Abcam, Cambridge, UK) and HRP-conjugated secondary antibodies (1:5000, Santa Cruz, Dallas, USA.). The blots were then visualized using the SuperSignal™ West Pico (Thermo Fischer) enhanced chemiluminescence detection system and recorded using a Gel Documentation 2000 system (Bio-Rad).

#### CXCL12 Bioactivity

*In vitro* CXCL12 bioactivity after its encapsulation in Ch/DS NC was evaluated by Jurkat migration assay (Zamproni, Mundim et al. 2017) .CXCL12 Ch/DS NC were added to 700 µL of migration buffer (RPMI supplemented with 0.5% bovine fetal serum) and were placed in 24 well plate. A Transwell cell insert (8 µm pore, Corning, New York, USA) was placed in each well containing the NC. Then, 4×10^5^ Jurkat cells (100 µL) were added in the apical chamber of the inserts. Migration buffer alone was used as negative control and a CXCL12 standard curve (50 ng/mL to 200 ng/mL) was used as a positive control. Cells were incubated at 37°C for 4 h in a 5% CO_2_ incubator. Transwell inserts were then removed and the number of cells that migrated in the lower chamber were measured by counting them with a Neubauer chamber.

#### NC internalization

Fluorescent Ch/DS NC were produced using a conjugated FITC-dextran-sulfate (500 kDa, Sigma). Four thousand N2a cells were seeded in the bottom of a 24-well plate and allowed to adhere overnight. Cells were then incubated with Ch/DS NC for 1, 3 or 24 hours and fixed with 4% paraformaldehyde. Nuclei were stained with DAPI (4′,6-diamidino-2-phenylindole, 1:500, Molecular Probes, Eugene, USA) and images were obtained using a scanning confocal inverted microscope (TCS, SP8 Confocal Microscope, Leica, Wetzlar, Germany). Fluorescence was calculated by dividing FITC integrated density by DAPI integrated density.

#### Effects of CXCL12 Ch/DS NC on normal brain cortex

Seven-week C57BL/6 female mice were submitted to surgery under anesthesia with intraperitoneal administration of acepromazine (1 mg/Kg, Vetnil, Louveira, Brazil), xylazine (10 mg/kg, Ceva, São Paulo, Brazil) and ketamine (100 mg/kg, Syntec, São Paulo, Brazil). Peroperative analgesia was provided by fentanyl (0.05 mg/Kg, Cristália, São Paulo, Brazil). All procedures were approved by the Committee of Ethics in the Use of Animals (CEUA/Unifesp 9745220217). Intracerebroventricular injection was performed according to a previously described protocol (DeVos and Miller 2013). Briefly, animals were placed under a stereotaxic frame, skin and skull were open. A 5 µL Hamilton syringe was used for injection (stereotaxic coordinates from bregma: AP +0.3 mm; ML +1.0 mm; DV −3.0 mm). Animals received 5 µL of PBS, Ch/DS NC, CXCL12 (20 ng/µL) or CXCL12 Ch/DS NC (20 ng CXCL12/ µL). Brains were collected 1 or 3 days later. Mice were anesthetized and euthanized by cervical dislocation. Ipsilateral brain cortex was immediately separated and frozen. Anterior ipsilateral brain cortex to injection was used for protein extraction and posterior ipsilateral brain cortex was used to RNA extraction.

#### Protein extraction

Frozen cortexes were homogenized in radioimmunoprecipitation assay (RIPA) buffer containing a protein inhibitor cocktail (Sigma). Samples were centrifuged at 7000 g for 5 min at 4°C. Supernatants were collected and frozen at −80 °C until further analysis. The protein amount in each sample was determined by Bio Rad DC Protein Assay (Bio Rad).

#### ELISA (Enzyme-Linked Immunosorbent Assay)

The amount of CXCL12, interleukin 6 (IL-6), interleukin 10 (IL-10) and tumor necrosis factor α (TNF-α) in protein homogenates from brain cortexes was measured by sandwich ELISA following the instructions of each detection Kit. Ninety mg of protein homogenates were used for CXCL12 quantification and 50 mg for IL-6, IL-10 and TNF-α quantification. The following kits were used: Mouse CXCL12 DuoSet ELISA (R&D systems, Minneapolis, USA), Mouse IL-6 ELISA Ready-SET-Go!^®^ (Thermo Fischer), Mouse IL-10 DuoSet ELISA (R&D systems) and Mouse TNF ELISA Set II (BD OptEIA™, BD Biosciences, San Diego, CA). The optical density was measured by an ELISA reader (EnSpire, Multimode Plate Reader, PerkinElmer do Brasil, São Paulo, Brazil).

#### RNA extraction and Quantitative PCR (qPCR)

Total RNA was isolated from brain cortex using Trizol® (Life Technologies, Thermo Fisher). RNA concentrations were determined using a NanoDrop ND-1000 instrument (Thermo Fisher). Reverse-transcriptase reactions were performed with the ImProm-II Reverse Transcription System (Promega, Madison, USA) using 2 μg total RNA. qPCR was performed using Brilliant® II SYBR® Green QPCR Master Mix (Applied Biosystems, Thermo Fisher) and the Mx3000P QPCR System; MxPro qPCR software was used for the analysis (Stratagene, San Diego, CA USA). Primers sequences are shown in Table 1. For quantification, target genes were normalized using Beta-actin (*Actb*) as housekeeping gene. The threshold cycles (Ct) were determined for each sample. The relative expression of mRNA was calculated using the 2^−ΔΔCt^ method (Livak and Schmittgen 2001) expressed relative to those from the PBS group.

**Table 1:**
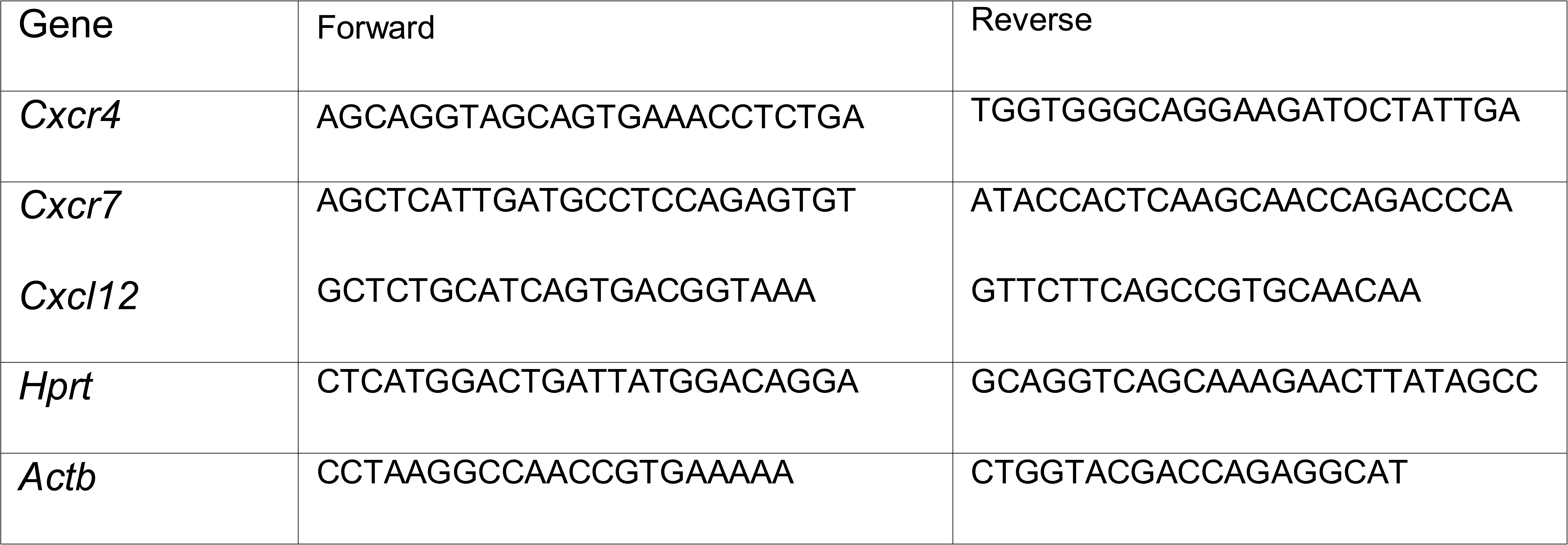
Primers sequences.

#### Stroke model

Cortical photothrombosis was induced by the Rose Bengal technique (Labat-gest and Tomasi 2013). Seven-week C57BL/6 female mice were submitted to surgery as described above. Anaesthetized mice were placed in a stereotaxic frame and the scalp incised, Rose Bengal (10mg/Kg, 15 mg/ml solution in sterile PBS) was injected intraperitoneally and the skull was illuminated for 20 min.

Immediately after the procedure, intracerebroventricular injection of 5 µL of PBS, NC Ch/DS, CXCL12 20 ng/mL or NC Ch/DS/CXCL12 20 ng/mL was performed as previously described. The scalp was sutured and the mice left to recover in a 30°C chamber for 1 h prior to returning to their home cage.

Mice were provided a dish of moist food in their home cage for the duration of their post-stroke survival. Mice were weighted every day. Brains were collected 3 days later. Mice were anesthetized and euthanized by cervical dislocation. Next, the brains were removed, stained with 2% 2,3,5-Triphenyl-tetrazolium (TCC, Alphatec, São José dos Pinhais, Brazil) at room temperature for 20 min and immersed in 4% paraformaldehyde for 24 h. To estimate the amount of cerebral tissue lost after stroke, the brains were sectioned into 2 mm coronal slices using a manual sectioning block, and the lost tissue area in each slice was determined by subtracting the area of the injured hemisphere from the area of the normal hemisphere using Image J software. The volume of brain tissue lost was determined by the sum of each slice area multiplied by the thickness (2 mm): lost area = Σ (area of contralateral side – area of ipsilateral side) x 2 as described (Llovera, Roth et al. 2014).

A second stroke model, permanent distal middle cerebral artery occlusion was performed as previously described (Llovera, Roth et al. 2014). A 1 cm skin incision between the ear and eye was made and the temporal muscle was separated. The middle cerebral artery (MCA) was identified at the skull, in the rostral part of the temporal area, dorsal to the retro-orbital sinus. The bone was thinned out with a drill right above the MCA branch and carefully withdraw. MCA was coagulated with monopolar electrosurgery (MedCir, São Paulo, Brazil). The temporal muscle was relocated to its position, covering the burr hole and skin sutured. Immediately after the procedure, intracerebroventricular injection of 5 µL of PBS, NC Ch/DS, CXCL12 20 ng/mL or NC Ch/DS/CXCL12 20 ng/mL was performed. The mice left to recover in a 30°C chamber for 1 h prior to returning to their home cage. Brains were collected 7 days later.

#### Behavioral assessment: Grid walking

To verify limb sensitivity, mice were recorded while walking over a grid for 4 min. The brain injury was in the left cortex, so as the mouse walked over the grid, its right paws falls, because it could not grasp the grid properly. The number of falls reflects the severity of the lesion (López-Valdés, Clarkson et al. 2014).

#### Ch/DS NC neurotoxicity

MTT (3-(4,5-dimethylthiazol-yl)2,5-diphenyltetrazolium bromide, Sigma) assay was performed to evaluate Ch/DS NC effect on N2a and NSC viability. Twenty thousand cells were grown overnight in 96-well plate and incubated for 24 h with different Ch/DS NC and CXCL12 Ch/DS NC concentrations (from 5 µL/mL to 100 µL/mL). In another test, N2a cells were incubated with 5 µL/mL or 50 µL/mL Ch/DS NC and CXCL12 Ch/DS NC for 4 or 24 hours. After that time, plate was washed with PBS, 3 times and incubated with regular media for 3 days. The MTT solution diluted in culture medium (1/10; 5 mg/L) was incubated for 2 h at 37 °C with treated cells. The MTT solution was removed and the formazan was dissolved in 100 μL dimethyl sulfoxide (DMSO, MP Biomedicals, Solon, USA). The plate was shaken for 15 min and optical density was read at 540 nm on an ELISA plate reader (Labsystems Multiskan MS, Helsinki, Finland).

#### Measurement of ROS Production

Reactive oxidative species (ROS) production was measured by using 2’,7’ – dichlorofluorescin diacetate (DCFDA, Sigma). DCFDA is cell-permeable and, once inside the cell, is de-esterified and turns to highly fluorescent 2′,7′-dichlorofluorescein upon oxidation. Briefly, N2a cells were harvested and seeded at 2×10^4^ cells per well on a black 96-well plate. Cells were allowed to attach overnight and were stained by incubation with 25 µM DCFDA in DMEM without phenol red for 45 min at 37°C. After washing with PBS, the cells were treated with Ch/DS NC, CXCL12 Ch/DS NC (from 5 µL/mL to 100 µL/mL) or H_2_O_2_ (positive control) and fluorescence/ROS was measured after 4 and 24 hours (Ex/Em: 485/535, RF-5301 Spectrofluorophotometer, Shimadzu, Kyoto, Japan).

#### Neurite Outgrowth

To evaluate impact of the CXCL12 Ch/DS NC on neuronal differentiation, N2a cells were plated in 24-well coverslips coated with 10 μg/mL of poly-L-lysine (Sigma) and 50 μg/mL of laminin (Sigma) and incubated in differentiation medium (high glucose DMEM, 0.5% FBS, 10 mM retinoic acid) with different Ch/DS NC and CXCL12 Ch/DS NC concentrations (from 5 µL/mL to 100 µL/mL). After 24 h, cells were fixed with 4 % paraformaldehyde and stained for β3-tubulin (anti-mouse IgG, 1:500, Sigma followed by Alexa 488 anti-mouse, 1:500, ThermoFisher Scientific and DAPI). Similarly, 4×10^4^ NSCs were seeded in laminin coated coverslips and incubated in EGF and FGF free medium with different Ch/DS NC concentrations (from 5 µL/mL to 100 µL/mL) for 24 hours. After 24 h, cells were fixed with 4 % paraformaldehyde and stained for doublecortin (guinea pig anti-DCX IgG, 1:400, Merck Millipore followed by Alexa 488 anti-mouse, 1:500, ThermoFisher Scientific and DAPI). Images were obtained using a scanning confocal inverted microscope (TCS, SP8 Confocal Microscope, Leica, Wetzlar, Germany). Neurite length was defined as the distance between the center of the cell soma and the tip of its longest neurite and was measured with Neurite Tracer (Image J).

#### Neurosphere migration

Neurospheres were plated in laminin coated coverslips, incubated with different Ch/DS NC concentrations (from 5 µL/mL to 100 µL/mL) for 24 h and were photographed using an inverted microscope (Olympus, Tokyo, Japan). The distance from the edge of the sphere to the farthest migrated cells was measured at four distinct locations per sphere using ImageJ and the average of this value was calculated (Kong, Fan et al. 2008).

#### Statistical analysis

Statistically significant differences were evaluated by one-way ANOVA followed by Tukey post-test using GraphPad Prism software version 5.01 (GraphPad Software, USA). Results are expressed as mean ± standard error of mean and were considered significant if p<0.05. The presence of the symbol * represents that data is different from control.

## Results

### Characterization of CXCL12 Ch/DS NC

Ch/DS NC and CXCL12 Ch/DS NC had a mean size of 186 and 143 nm, respectively (Table 2). Their zeta potential was in average −25 mV and EE was 75% while CXCL12 loading was 19 ng per stock µL formulation.

**Table 2:** NC characterization.

|  | Ch/DS NC | CXCL12 Ch/DS NC |
| --- | --- | --- |
| Size (nm) | 186,4 ± 76,1 | 142,7 ± 56,9 |
| Zeta potential (mV) | -25,36 ± 0,73 | -24,10 ± 1,5 |
| EE (%) | n.a. | 75,12 ± 4,58 |
| DL (ng/μL) | n.a. | 18,78 ± 1,13 |

n.a: not applicable

Based on CXCL12 loading, 2.5 µL and 5 µL of CXCL12 Ch/DS NC stock solution containing 50 ng and 100 ng of CXCL12, respectively, were used for the following experiments.

We found minimal CXCL12 release from CXCL12 Ch/DS NC *in vitro*, with only 0.5% of the total amount of protein released after 7 days (Figure 2A), although we confirmed the presence of CXCL12 in the pellet by Western blot (Figure 2B). CXCL12 bioactivity was preserved in the complexes and calculated as 80,4% ± 9,1% of free CXCL12 (n=3) (Figure 2C).

**Figure 1:**
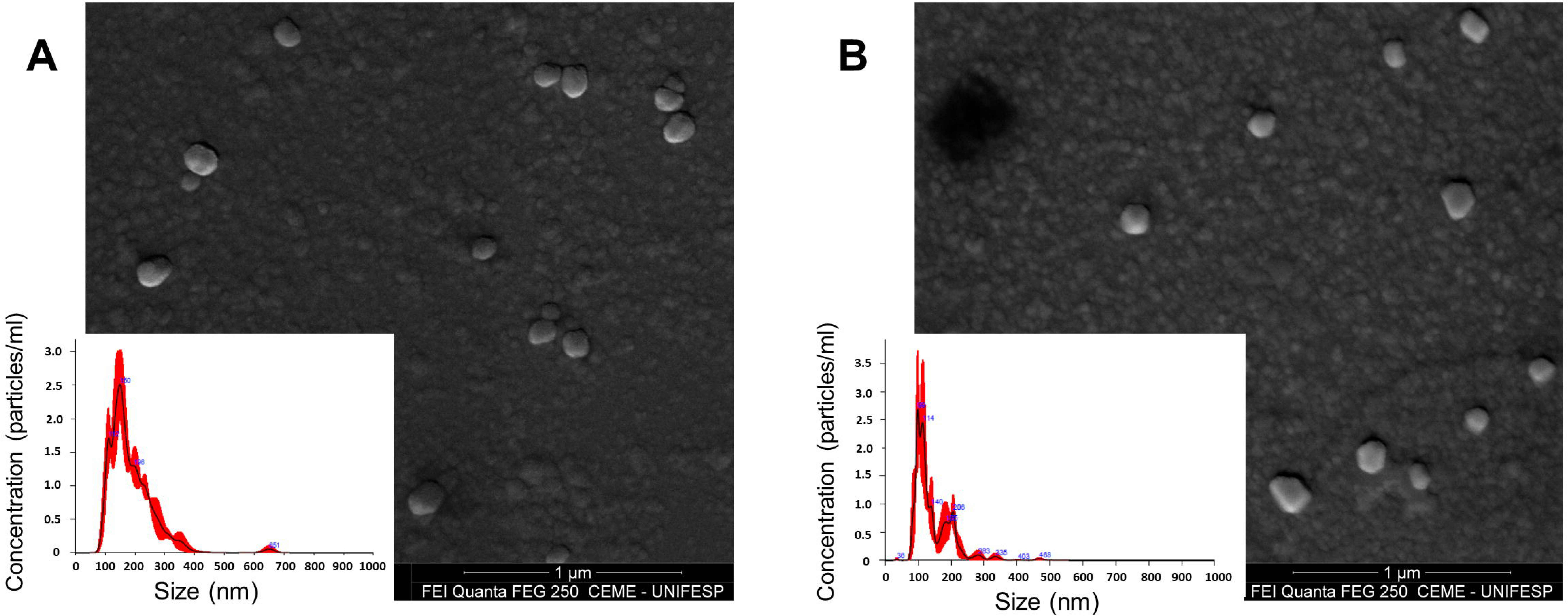
NC characterization. Size distribution graph from Nanosight (Malvern) and SEM pictures for (A) Ch/DS NC and (B) CXCL12 Ch/DS NC. NC: nanocomplexes; Ch: chitosan; DS: dextran sulfate; SEM: scanning electron microscopy.

**Figure 2:**
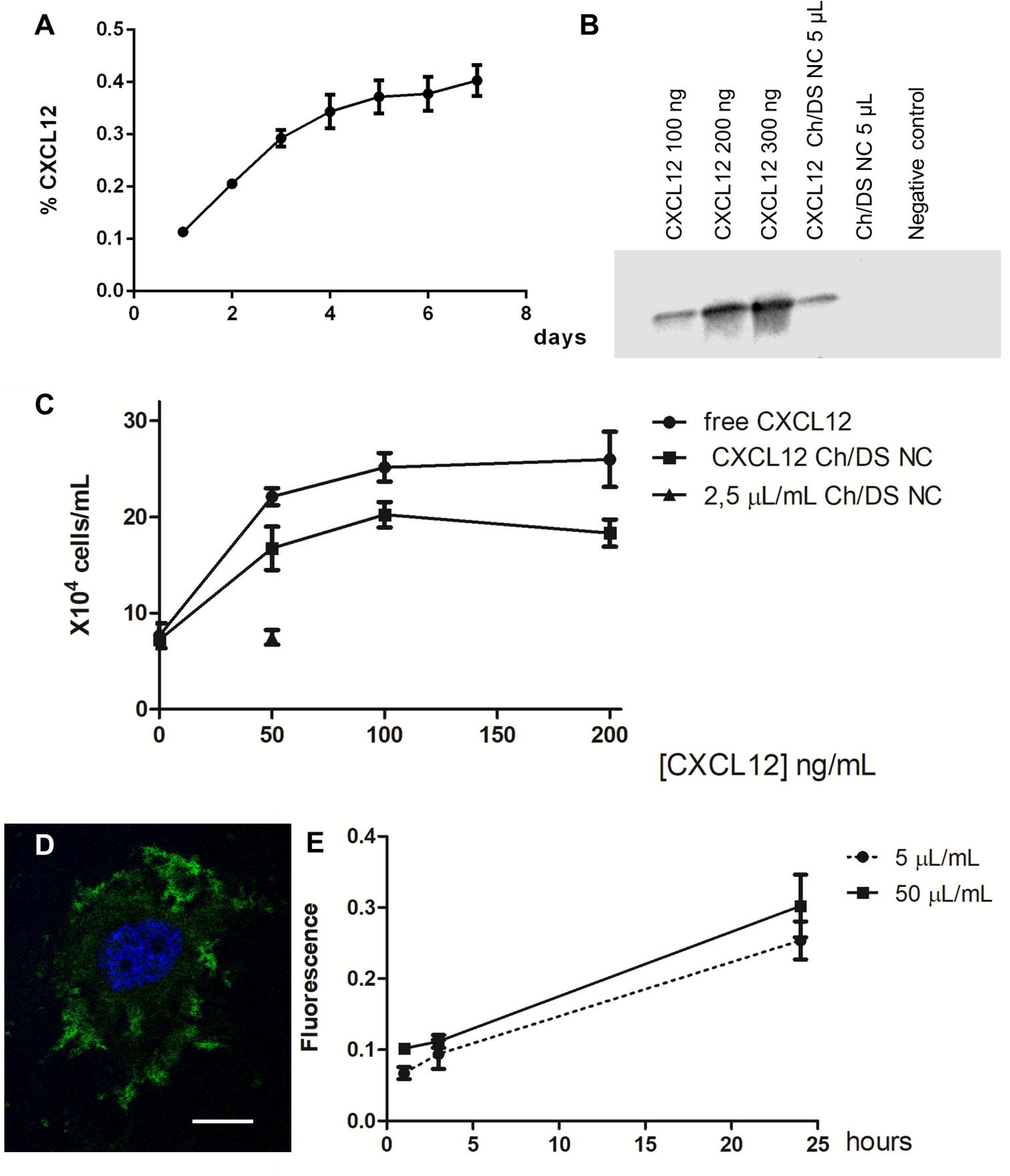
NC characterization. (A) Percentage of CXCL12 incorporated in Ch/DS NC released in 7 days *in vitro*. Less than 0.5% was released (n=3). (B) Detection of CXCL12 incorporated in NC by Western Blotting (n=2). (C) Jurkat transwell migration. Bioactivity of CXCL12 in Ch/DS NC was 80.4% ± 9.1% of free CXCL12 (n=3). (D) Representative figure showing fluorescent Ch/DS NC cell internalization (Scale bar = 20 µm). (E) Fluorescent Ch/Ds NC cell internalization dynamics (n=3 high field images evaluated per condition). Data expressed as mean ± standard error of mean. NC: nanocomplexes; Ch: chitosan; DS: dextran sulfate.

N2a cells were able to internalize NC. Internalization was shown to increase with time, but not with increasing NC concentration in media (Figures 2D and E).

### CXCL12 Ch/DS NC were capable of increase CXCL12 receptors in vivo

First, we assessed whether CXCL12 Ch/DS NC were able to increase the concentration of CXCL12 in the cerebral cortex or its receptors after intracerebroventricular injection. CXCL12 protein levels were quantified in the cortex ipsilateral to the injection. No increased CXCL12 concentration was detected in any of the groups who received free CXCL12 or CXCL12 Ch/DS NC at day 1 and 3 (Figure 3A). Similarly, no increased of *Cxcl12* gene expression was observed in any of the groups (Figure 3B). However, the gene expression of *Cxcr7*, a CXCL12 receptor, significantly increased 1 day after injection of CXCL12 Ch/DS NC compared to PBS (Figure 3C). *Cxcr4* gene expression, another receptor for CXCL12, significantly increased 3 days after injection of CXCL12 Ch/DS NC compared to PBS (Figure 3D).

**Figure 3:**
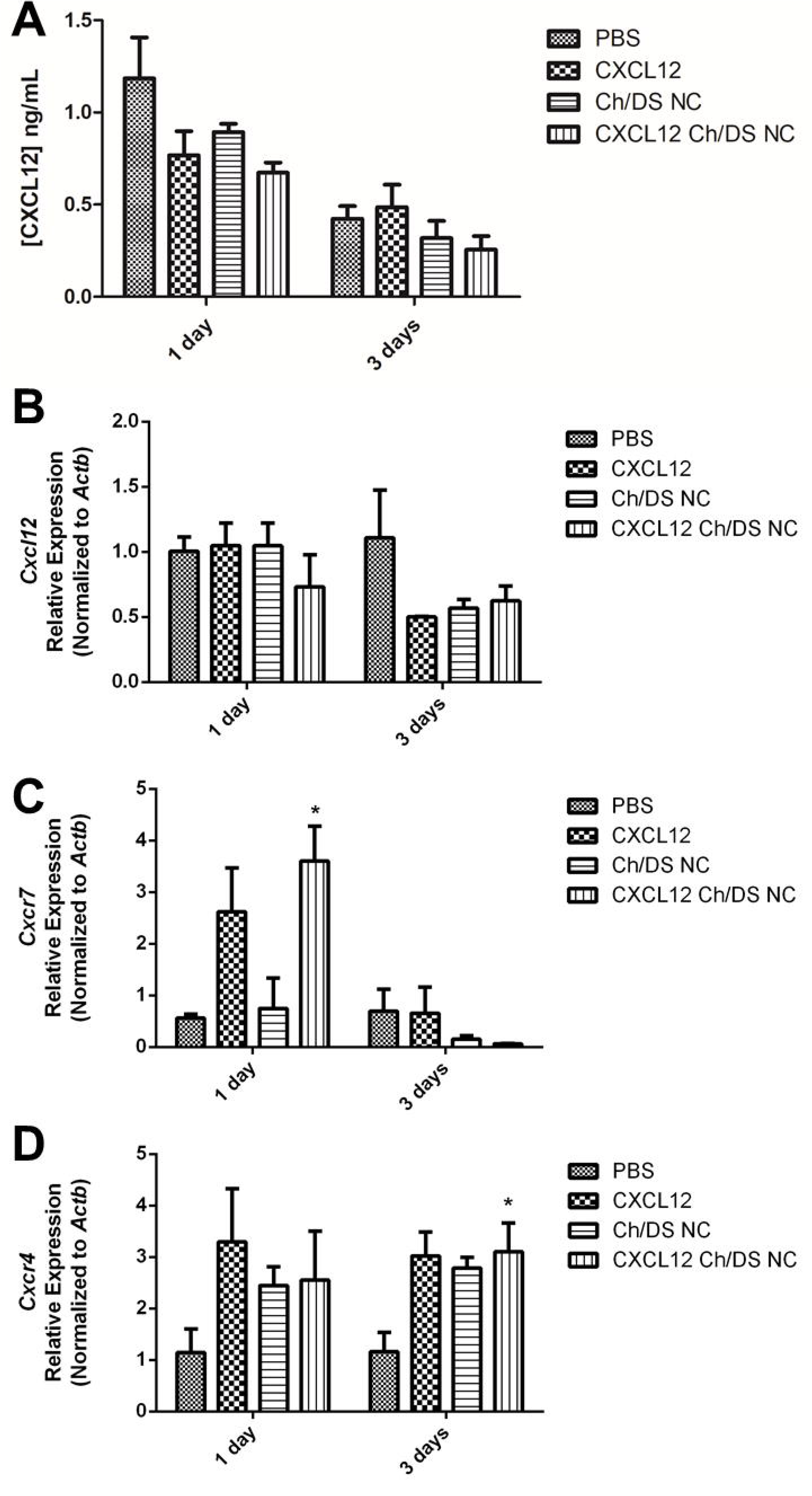
CXCL12 Ch/DS NC were capable to increase CXCL12 receptors *in vivo*. (A) CXCL12 concentration (ng/mL) in the cortex measured by ELISA (n=3). (B) *Cxcl12* relative expression in the cortex (n=3). (C) *Cxcr7* relative expression in the cortex (*= p < 0.05, one-way Anova plus Tukey posttest, n=3). (D) *Cxcr4* relative expression in the cortex (*= p < 0.05, one-way Anova plus Tukey posttest, n=3). Data expressed as mean ± standard error of mean. NC: nanocomplexes; Ch: chitosan; DS: dextran sulfate; ELISA: Enzyme-Linked Immunosorbent Assay, *Actb*: beta-actin.

### Ch/DS NC are deleterious in stroke

After characterization, we attempted to verify the CXCL12 Ch/DS NC therapeutic ability in a photothrombotic mice stroke model. Surprisingly, animals who received empty Ch/DS NC presented increased infarct area (Figures 4A and B) associated with worse performance in the grid test (Figure 4C). To confirm these data the experiment was repeated using a different model of stroke, the permanent distal occlusion of the left middle cerebral artery. Again, a significant increase in ischemic injury was observed in animals receiving Ch/DS NC (Figure 5).

**Figure 4:**
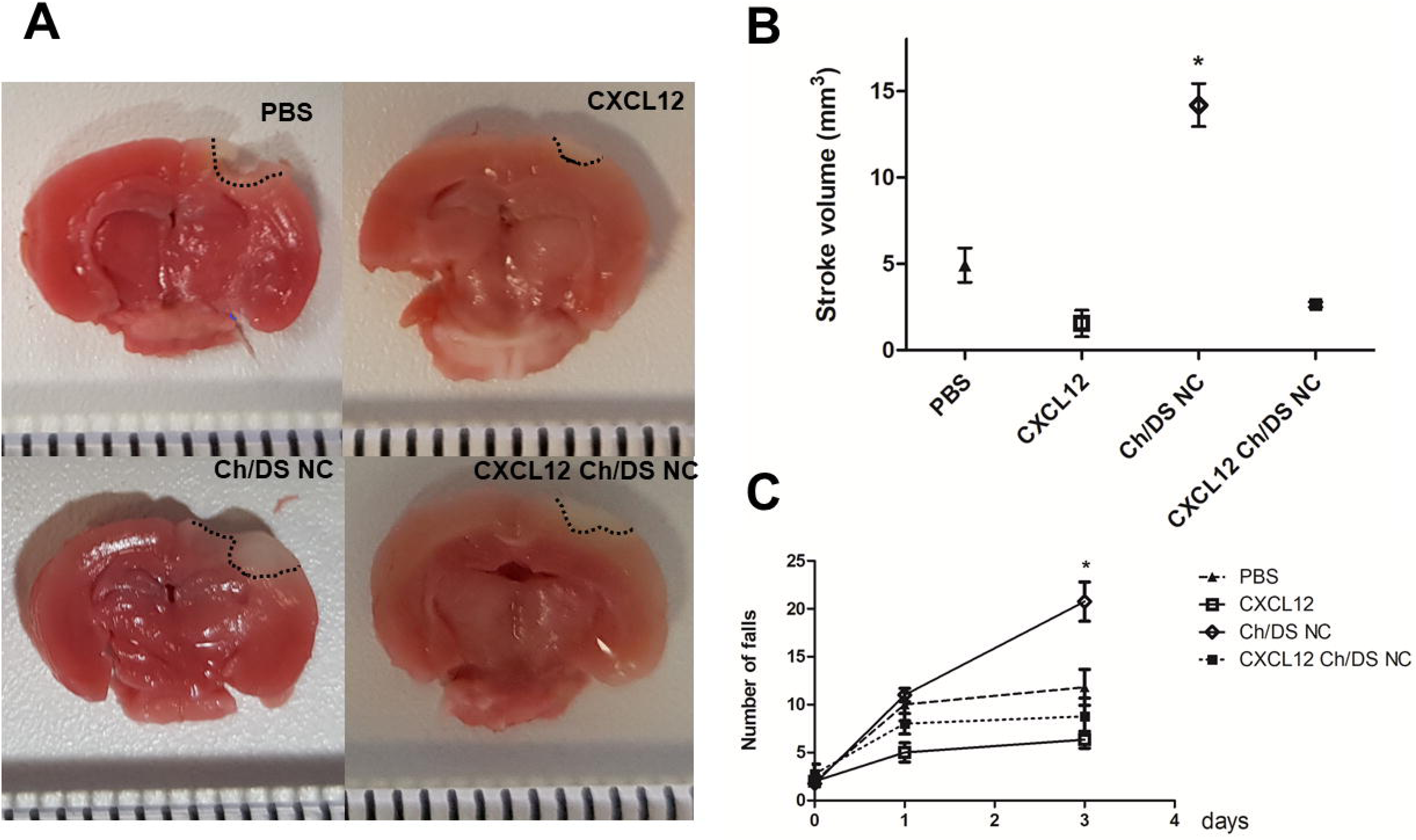
Ch/DS NC were deleterious in cortical photothrombosis stroke. (A) TTC staining of brain slices. Unstained area corresponds to death tissue. (B) Quantification of stroke area by TTC method (*= p < 0.001, one-way Anova plus Tukey posttest n=5 group PBS, n= 3 group CXCL12, n= 4 groups Ch/DS NC and CXCL12 Ch/DS NC). (C) Number of paws falls in the grid walking test in mice (*= p = 0.001, one-way Anova plus Tukey posttest n=5 group PBS, n= 3 group CXCL12, n= 4 groups Ch/DS NC and CXCL12 Ch/DS NC). Data expressed as mean ± standard error of mean. NC: nanocomplexes; Ch: chitosan; DS: dextran sulfate; TTC: 2,3,5-triphenyltetrazolium chloride.

**Figure 5:**
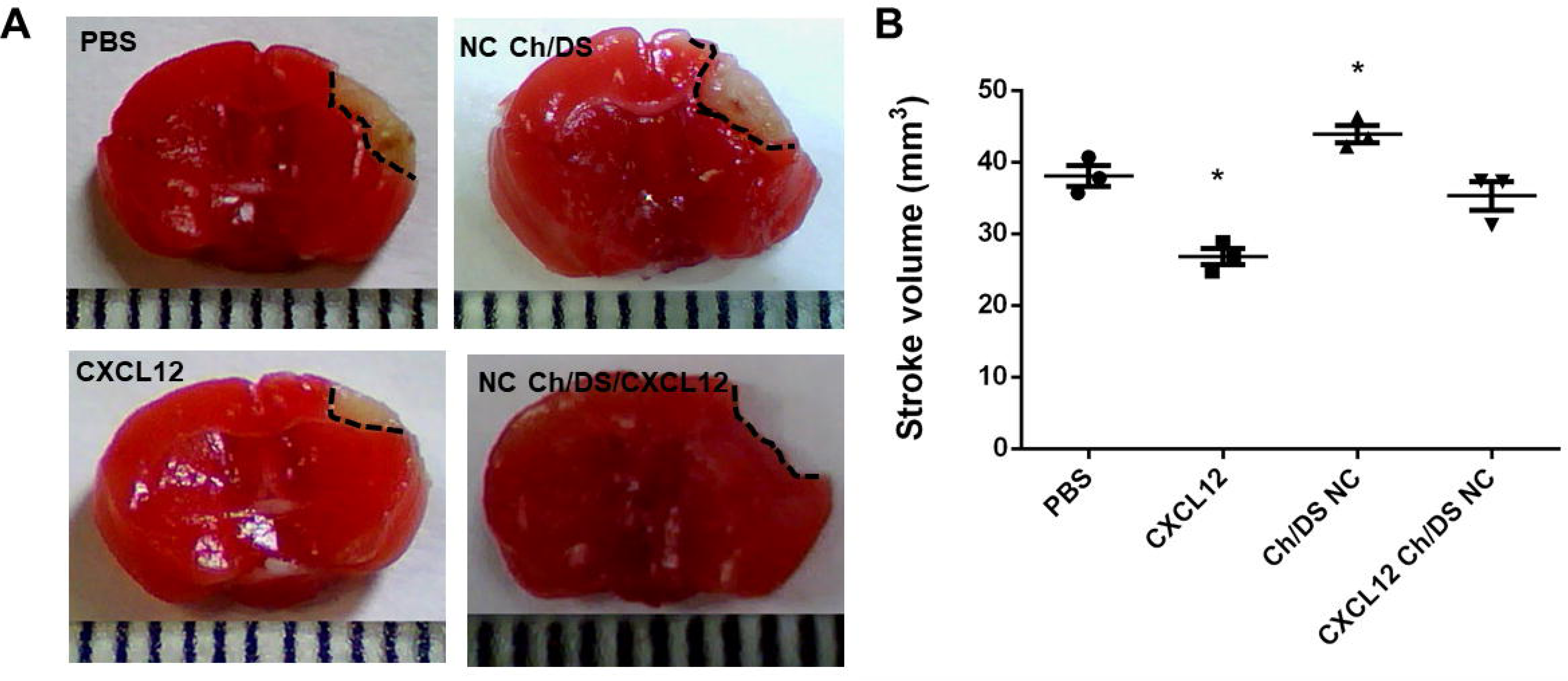
Ch/DS NC were deleterious in middle cerebral artery occlusion stroke model (A) TTC staining of brain slices. Unstained area corresponds to death tissue. (E) Quantification of stroke area by TTC method (*= p < 0.05, one-way Anova plus Tukey posttest, n=3). Data expressed as mean ± standard error of mean. NC: nanocomplexes; Ch: chitosan; DS: dextran sulfate; TTC: 2,3,5-triphenyltetrazolium chloride.

### Ch/DS NC decreases cell viability

The increase in the ischemic area in animals that received the Ch/DS NC brought us the concern about their safety. So, we conducted assays to verify the compatibility of NC with brain tissue.

First, cell viability was verified by MTT assay, using two cellular types, a murine neuroblastoma cell lineage (N2a) and a primary culture of neural stem cells (NSCs) extracted from SVZ, for 24 h. We observed that in both N2a and NSCs viability was slightly decreased after exposure to Ch/DS NC for 24 h (Figures 6A and B). Viability was maintained when CXCL12 was added to the formulation (Figure 6C). Since oxidative stress is one of the causes of death in cells exposed to nanoparticles (Khanna, Ong et al. 2015), we verified whether NC could increase ROS production using DCFDA. We found that neither Ch/DS NC or CXCL12 Ch/DS NC increased ROS production in 4 or 24 h (Figure 6D and E).

**Figure 6:**
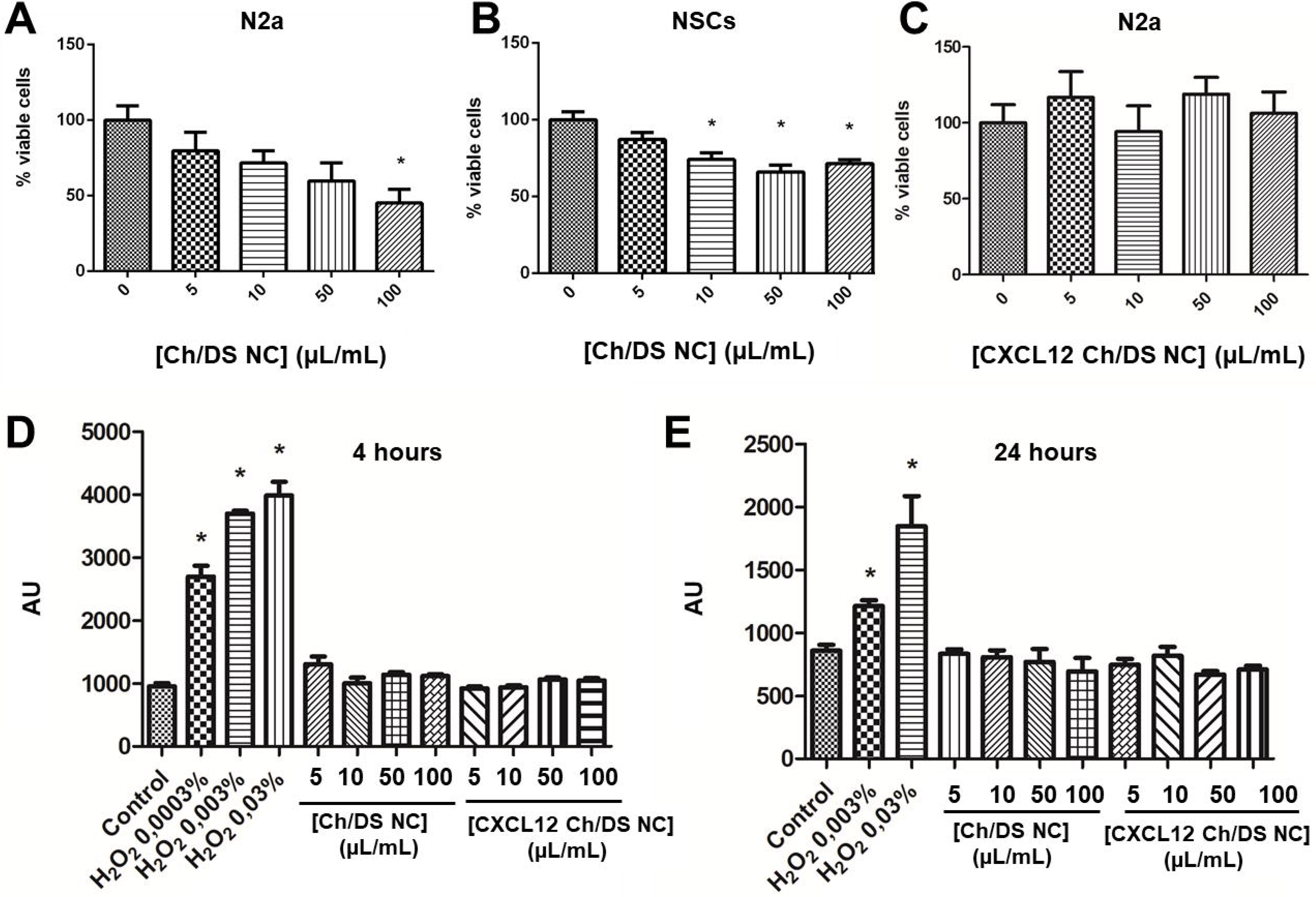
Neurotoxicology screening for Ch/DS NC. (A) Viability of N2a cells by MTT after 24 h exposed to different concentrations of Ch/DS NC (µL/mL) (*= p < 0.01, one-way Anova plus Tukey posttest, n= 6). (B) Viability of NSCs by MTT after 24 h exposed to different concentrations of Ch/DS NC (µL/mL) (*= p < 0.0001, one-way Anova plus Tukey posttest, n= 5). (C) Viability of N2a cells by MTT after 24 h exposed to different concentrations of CXCL12 Ch/DS NC (µL/mL) (one-way Anova plus Tukey posttest, n= 6). (D) ROS production by DCFDA fluorescent assay in 4 hours (*= p < 0.0001, one-way Anova plus Tukey posttest, n= 6). (E) ROS production by DCFDA fluorescent assay in 24 hours (*= p < 0.0001, one-way Anova plus Tukey posttest, n= 6). Data expressed as mean ± standard error of mean. NC: nanocomplexes; Ch: chitosan; DS: dextran sulfate; NSCs: neural stem cells; MTT: 3-(4,5-dimethylthiazol-yl)2,5-diphenyltetrazolium bromide; DCFDA: 2’,7’ –dichlorofluorescin diacetate; AU: arbitrary units.

However, during the experiments we realized that when trying to incubate the cells with NC for longer periods, at the end of 72 hours there were practically no more cells in the wells. To confirm this, we repeated the MTT assay, this time exposing the cells to the NCs for 4 or 24 hours. After this period the NC containing medium was removed, the cells washed and incubated with normal culture medium (Figure 7A). At 72 hours, the MTT assay confirmed there were no more viable cells in the wells (Figure 7B and C). Two possibilities occurred to us to explain this result: either the cells died from exposure or were stopping to divide. For this reason, we decided to check for a possible interaction of NCs with cytoskeleton.

**Figure 7:**
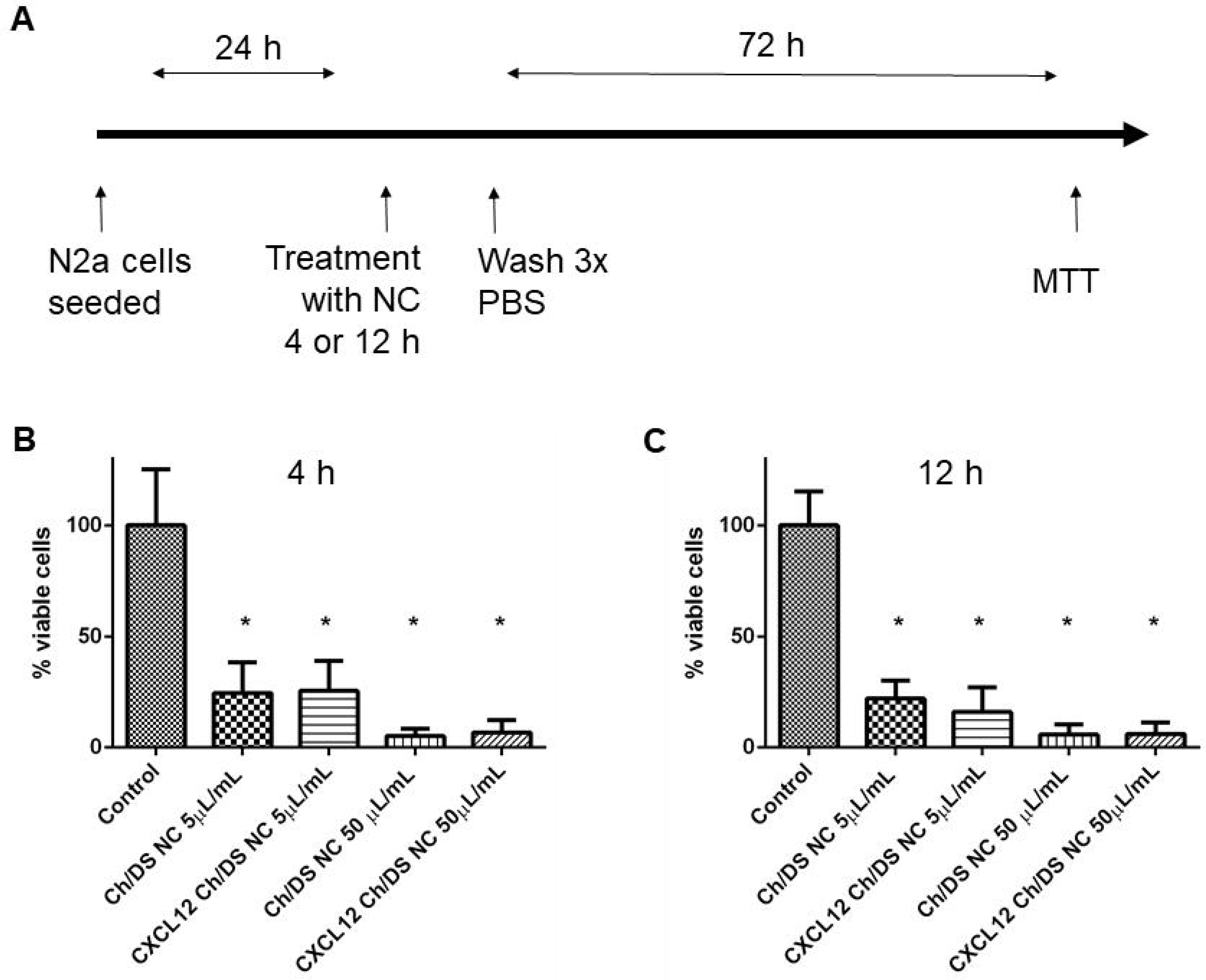
Ch/DS NC reduces N2a viability in 72 hours. (A) Schematic representation of the experiment protocol. Cells were exposed to NC for 4 ou 24 hours, then washed and incubated with regular media till MTT assay in 72 hours. (B) Viability of N2a by MTT in 72 h after 4 h exposed to Ch/DS NC or CXCL12 Ch/DS NC (5 or 50 µL/mL) (*= p < 0.0001, one-way Anova plus Tukey posttest, n= 8). (C) Viability of N2a by MTT in 72 h after 24 h exposed to Ch/DS NC or CXCL12 Ch/DS NC (5 or 50 µL/mL) (*= p < 0.0001, one-way Anova plus Tukey posttest, n= 8). Data expressed as mean ± standard error of mean. NC: nanocomplexes; Ch: chitosan; DS: dextran sulfate; MTT: 3-(4,5-dimethylthiazol-yl)2,5-diphenyltetrazolium bromide; PBS: phosphate buffer saline.

### Ch/DS NC decreases neurite length

To evaluate the interaction of NCs with cytoskeleton, we evaluated the morphology of neural cells and NSC ability to migrate. Neurite length in N2a and NSCs was dramatically decreased by Ch/DS NC (Figure 8A). Also, Ch/DS NC impaired neuroblasts emigration from neurospheres (Figure 8A). When CXCL12 was added to the formulation, neurite outgrowth impairment was not significant, but their size remained 50 to 80 µm shorter (Figure 8B).

**Figure 8:**
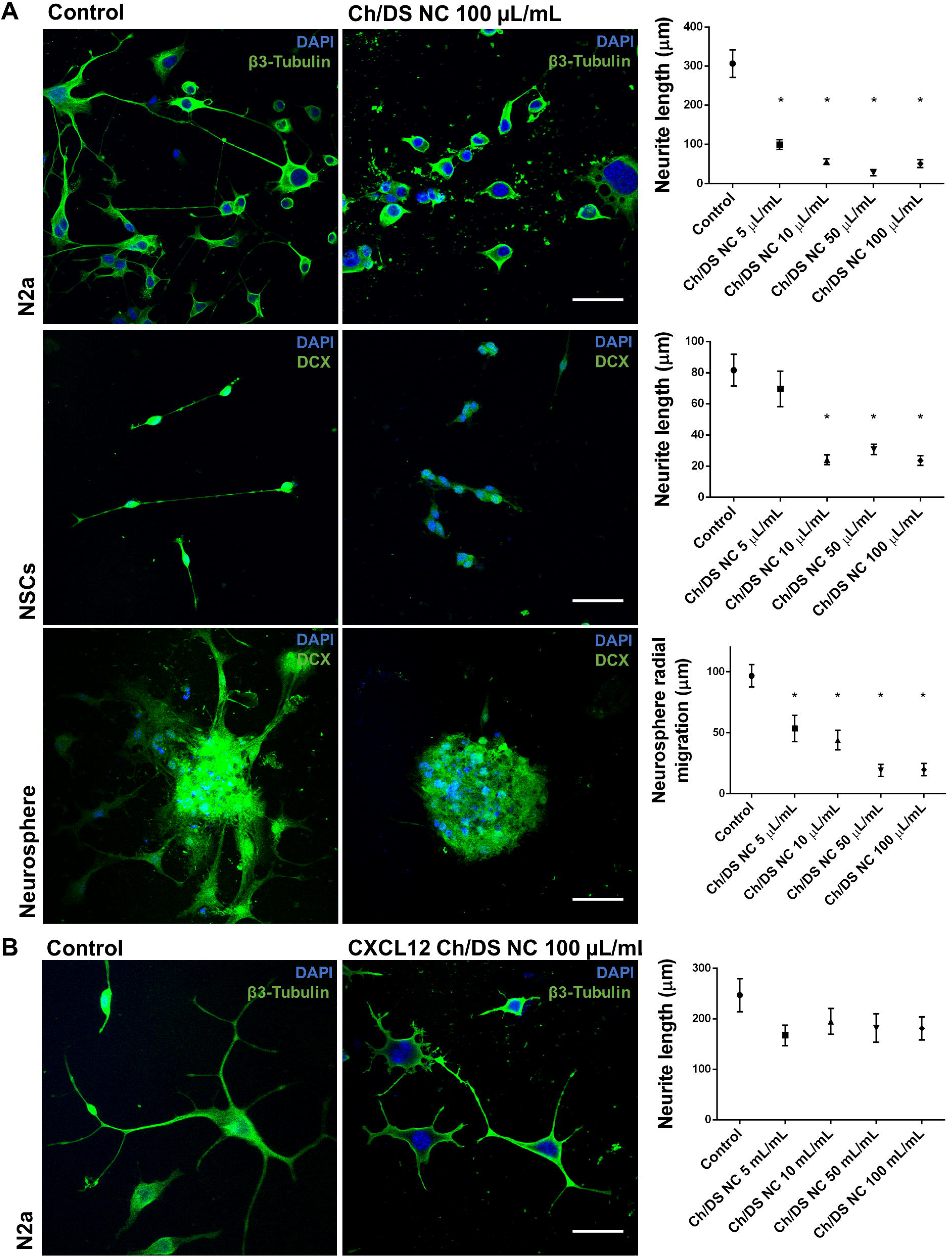
Ch/DS NC decreases neurite length. (A) Immunocytochemistry of N2a cells, NSCs and neurospheres incubated with Ch/DS NC (left panels, Scale bar = 50 µm) and graphical re-presentation of the numerical quantification of neurite length of N2a cells (superior right panel, *=p < 0.0001, n= 31, control group, n=32, Ch/DS NC 5 µL/mL group, n=30 Ch/DS NC 10 µL/mL group, n=23 Ch/DS NC 50 µL/mL group and Ch/DS NC 100 µL/mL group) and NSCs (middle right panel, *=p < 0.0001, n= 18, control group, n=26, Ch/DS NC 5 µL/mL group, n=22 Ch/DS NC 10 µL/mL group, n=21 Ch/DS NC 50 µL/mL group and n=13 Ch/DS NC 100 µL/mL group) and graphical representation of the numerical quantification of neurosphere migration (inferior right panel, *=p < 0.0001, n= 14, control group and Ch/DS NC 5 µL/mL group, n=16 Ch/DS NC 10 µL/mL group, n=13 Ch/DS NC 50 µL/mL group and n=11 Ch/DS NC 100 µL/mL group). (B) Immunocytochemistry of N2a cells incubated with CXCL12 Ch/DS NC (left panel, Scale bar = 50 µm) and graphical representation of the numerical quantification of neurite length (right panel, p > 0.05, n= 20). Data expressed as mean ± standard error of mean. NC: nanocomplexes; Ch: chitosan; DS: dextran sulfate; NSCs: neural stem cells; DAPI: 4′,6-diamidino-2-phenylindole; DCX: doublecortin.

### CXCL12 Ch/DS NC upregulates the production of anti-inflammatory interleukins in brain cortex

To detect a potential effect of CXCL12 Ch/DS NC on inflammation, IL-10, IL-6 and TNF-α concentrations in the ipsilateral normal cortex to the injection were quantified. Twenty-four hours after CXCL12 Ch/DS NC injection IL-6 concentration was increased in brain tissue when compared to PBS (Figure 9A). IL-10 production increased in animals treated with CXCL12 Ch/DS NC after 72 h (Figure 9B). No impact on TNF-α concentrations was observed disregard of treatments (Figure 9C).

**Figure 9:**
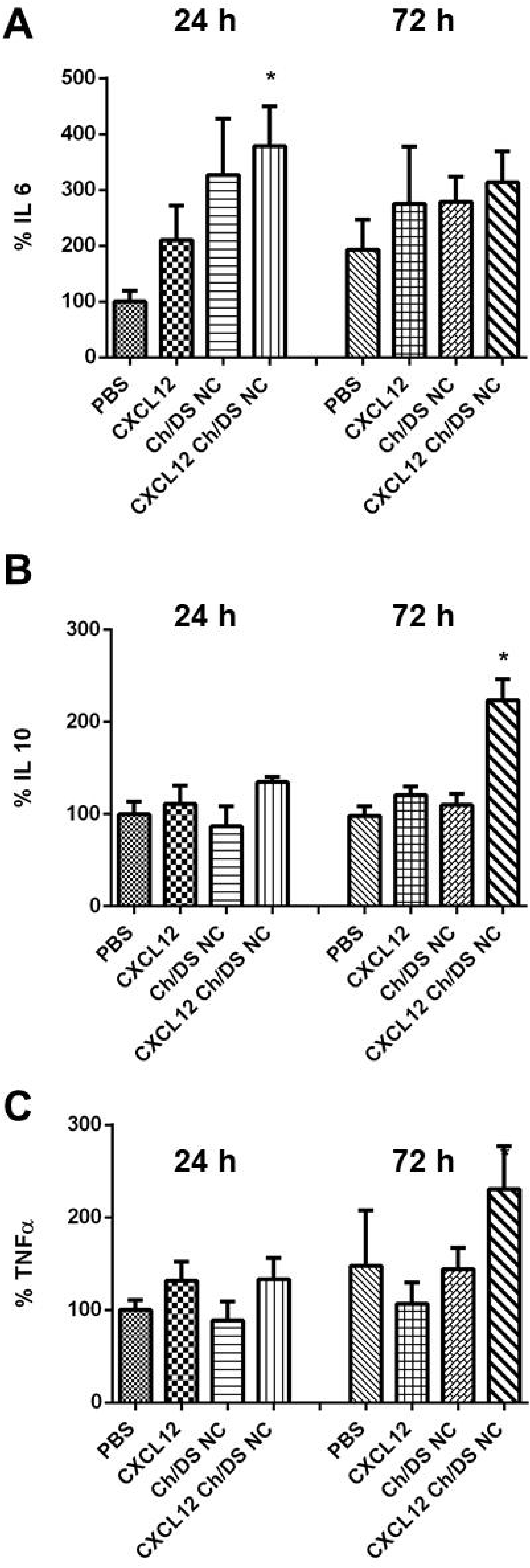
Inflammatory mediators on brain cortex. (A) IL-6 concentration (expressed in percentage of the control PBS 24 h group) in the cortex measured by ELISA (*= p < 0.05, one-way Anova plus Tukey posttest n=6). (B) IL-10 concentration (expressed in percentage of the control PBS 24 h group) in the cortex measured by ELISA (*= p < 0.001, one-way Anova plus Tukey posttest n=6). (C) TNF-α concentration (expressed in percentage of the control PBS 24 h group) in the cortex measured by ELISA (n=6). Data expressed as mean ± standard error of mean. NC: nanocomplexes; Ch: chitosan; DS: dextran sulfate; IL: interleukin; TNF: tumor necrosis factor.

## Discussion

In this study, we found evidence that Ch/DS NC exert deleterious effects on brain tissue when administered intracerebroventricularly in mice. Indeed, the experimental group that received Ch/DS NC showed increased stroke volume and presented worsened performance in the grid walking test. Also, a decrease in N2a and NSCs viability and neurite length *in vitro* was observed.

The Ch/DS NC used in this work are not new and have already been described previously (des Rieux, Ucakar et al. 2011, Yin, Bader et al. 2013, Bader, Li et al. 2015). The characterization conducted in this work show NC with characteristics very similar to those previously described (Bader, Li et al. 2015).

Reports have shown that engineered nanomaterials can enter the brain and cause tissue damage (Simkó and Mattsson 2014, Lima, Guimarães et al. 2019), although data regarding nanoparticles toxicity remain limited. In this work we chose to inject the complexes directly into the brain, since the dextran sulfate used weights 500 kDa and is known to not permeate the healthy blood-brain barrier.

Chitosan nanoparticles have already been shown to cause zebrafish embryos death and malformation (Hu, Qi et al. 2011) and more recently, chitosan-coated zein nanoparticles were shown to enhance anxiety and depression in rats (Lima, Guimarães et al. 2019). Although chitosan preparation involves basic purification methods like demineralization and deproteinization, chitosan may contain some impurities, such as ash, heavy metals, or proteins. The purity level of chitosan affects its biological properties like immunogenicity or biodegradability. Therefore, chitosan should be of high purity and be free of contaminants (including the level of endotoxins) (Szymańska and Winnicka 2015). The chitosan used in this work was highly pure following suppliers’ information.

Nanoparticles can exert deleterious effects by acting in several pathways. The primary mechanism of nanoparticle toxicity involves the production of ROS and free radicals leading to oxidative stress reactions and inflammation (Yuan, Hu et al. 2015, Teleanu, Chircov et al. 2018). In this study, we found no evidence for increased oxidative stress reactions.

Many studies have indicated that nanomaterial exposure induces cytoskeleton disruption and cell-morphology changes (Xu, Piett et al. 2013, Kang, Liu et al. 2016, Liu, Kang et al. 2017). In this study, we observed a change in β3-tubulin and doublecortin cell distribution after Ch/DS NC exposure with a dramatic decrease of neurite outgrowth. Besides, there was impairment of NSCs migration from neurospheres. Because all brain functions rely critically upon the normal development of neuronal structures and network circuitry, any perturbation of these processes will lead to central nervous system dysfunctionality (Xu, Piett et al. 2013).

The addition of CXCL12 to NC formulation was able to attenuate deleterious effects observed after Ch/NC NC exposure. Free CXCL12 has been already shown to increase viability measured by MTT assay in human colorectal cancer cells and in osteosarcoma cells which can justify the better MTT viability data when neuroblastoma cells where in presence of CXCL12 Ch/DS NC (Li, Yu et al. 2008, Liao, Fu et al. 2015).

CXCL12 forms a complex with CS and DS and it is possible that in its absence, some charges are left free to impact electrical neuronal activity by induction of depolarization in the neuronal membrane potential (Teleanu, Chircov et al. 2018). CXCL12 regulates axonal elongation and branching and has previously been shown to promote axonal growth even in the presence of potent chemorepellent molecules (Chalasani, Sabelko et al. 2003, Chalasani, Sabol et al. 2007). Treatment with CXCL12 was sufficient to overcome neurite outgrowth inhibition mediated by central nervous system myelin in cultured postnatal dorsal root ganglion neurons (Opatz, Küry et al. 2009). Also, CXCL12 Ch/DS NC modulated the inflammatory response beneficially by inducing IL-6 and IL-10 production in mice brain cortex. IL-6 is a key early mediator of the inflammatory and overall immune responses. Many different cells including astrocytes, mast cells, monocytes, macrophages, fibroblasts and endothelial cells can secrete IL-6 after cerebral ischemia (Li, Qi et al. 2017). IL-6 appears to have a biphasic, or even polyphasic, role in cerebral ischemia. Evidence for a neuroprotector role of IL-6 in cerebral ischemia is suggested by recent studies showing impaired neurogenesis in mice lacking IL-6 (Famakin 2014). IL-10 may protect against ischemic injury during the acute phase of stroke (Protti, Gagliardi et al. 2013). Expression of IL-10 is elevated during most central nervous system diseases and promotes survival of neurons and glial cells in the brain by blocking the effects of proapoptotic cytokines and by promoting expression of cell survival signals (Strle, Zhou et al. 2001). Neuroinflammatory modulation may represent one of the mechanisms by which CXCL12 Ch/DS NC treated mice performed better than Ch/DS NC treated mice after stroke.

CXCL12 could be delivered in its free form. Our data showed that free CXCL12 was able to reduce ischemic injury. However as previously attested, the chemokine in its free form is unstable and prone to inactivation. In this case, other formulations capable of encapsulating CXCL12 could be useful for chemokine delivery (Zamproni, Mundim et al. 2017).

Although not yet used in stroke therapy, nanoparticulate systems and in particular chitosan-based nanoparticles have been extensively studied as promising for treatment in a number of conditions. Unfortunately, few articles seek to investigate possible deleterious consequences of these complexes. The authors would like to draw attention to the need for a better understanding of the neurotoxicity of these compounds and the need to design new neuroprotective therapies.

## Conclusions

Here we evaluated Ch/DS NC as a possible carrier for CXCL12 in a mouse model of stroke. However, the complexes turned out to be neurotoxic. Our results add new data on nanoparticle neurotoxicity and may help to better understand the complex interactions of the nanostructures with biological components and to design new neuroprotective therapies.

## Acknowledgments

We kindly thank FAPESP (MP: 2012/00652-5; LNZ: 2013/16149-3; 2014/23797-4) and CNPq (LNZ: 380304/2017-1; MP: 402319/2013-3; 465656/2014-5) for financial support. We also thank the Eletronic Microcospy Center (CEME) of the Federal University o São Paulo for helping with SEM pictures and Professor Karin do Amaral Riske for the use of Zetasizer.

## References

Adelita, T., et al. (2017). “Proteolytic processed form of CXCL12 abolishes migration and induces apoptosis in neural stem cells in vitro.” Stem Cell Res 22: 61–69.

Bader, A. R., et al. (2015). “Preparation and characterization of SDF-1α-chitosan-dextran sulfate nanoparticles.” J Vis Exp(95): 52323.

Chalasani, S. H., et al. (2003). “A chemokine, SDF-1, reduces the effectiveness of multiple axonal repellents and is required for normal axon pathfinding.” J Neurosci 23(4): 1360–1371.

Chalasani, S. H., et al. (2007). “Stromal cell-derived factor-1 antagonizes slit/robo signaling in vivo.” J Neurosci 27(5): 973–980.

Cheng, X., et al. (2017). “Elevated Serum Levels of CXC Chemokine Ligand-12 Are Associated with Unfavorable Functional Outcome and Mortality at 6-Month Follow-up in Chinese Patients with Acute Ischemic Stroke.” Mol Neurobiol 54(2): 895–903.

des Rieux, A., et al. (2011). “3D systems delivering VEGF to promote angiogenesis for tissue engineering.” J Control Release 150(3): 272–278.

DeVos, S. L. and T. M. Miller (2013). “Direct intraventricular delivery of drugs to the rodent central nervous system.” J Vis Exp(75): e50326.

Famakin, B. M. (2014). “The Immune Response to Acute Focal Cerebral Ischemia and Associated Post-stroke Immunodepression: A Focused Review.” Aging Dis 5(5): 307–326.

Hu, Y. L., et al. (2011). “Toxicity evaluation of biodegradable chitosan nanoparticles using a zebrafish embryo model.” Int J Nanomedicine 6: 3351–3359.

Kang, Y., et al. (2016). “Potential Links between Cytoskeletal Disturbances and Electroneurophysiological Dysfunctions Induced in the Central Nervous System by Inorganic Nanoparticles.” Cell Physiol Biochem 40(6): 1487–1505.

Khanna, P., et al. (2015). “Nanotoxicity: An Interplay of Oxidative Stress, Inflammation and Cell Death.” Nanomaterials (Basel) 5(3): 1163–1180.

Kim, D. H., et al. (2015). “Enhancing neurogenesis and angiogenesis with target delivery of stromal cell derived factor-1α using a dual ionic pH-sensitive copolymer.” Biomaterials 61: 115–125.

Kong, H., et al. (2008). “AQP4 knockout impairs proliferation, migration and neuronal differentiation of adult neural stem cells.” J Cell Sci 121(Pt 24): 4029–4036.

Labat-gest, V. and S. Tomasi (2013). “Photothrombotic ischemia: a minimally invasive and reproducible photochemical cortical lesion model for mouse stroke studies.” J Vis Exp(76).

Li, J. K., et al. (2008). “Inhibition of CXCR4 activity with AMD3100 decreases invasion of human colorectal cancer cells in vitro.” World J Gastroenterol 14(15): 2308–2313.

Li, W. X., et al. (2017). “Different impairment of immune and inflammation functions in short and long-term after ischemic stroke.” Am J Transl Res 9(2): 736–745.

Li, Y., et al. (2015). “CXCL12 Gene Therapy Ameliorates Ischemia-Induced White Matter Injury in Mouse Brain.” Stem Cells Transl Med 4(10): 1122–1130.

Liao, Y. X., et al. (2015). “AMD3100 reduces CXCR4-mediated survival and metastasis of osteosarcoma by inhibiting JNK and Akt, but not p38 or Erk1/2, pathways in in vitro and mouse experiments.” Oncol Rep 34(1): 33–42.

Lima, V. S., et al. (2019). “Depression, anxiety-like behavior, and memory impairment in mice exposed to chitosan-coated zein nanoparticles.” Environ Sci Pollut Res Int.

Liu, J., et al. (2017). “Zinc oxide nanoparticles induce toxic responses in human neuroblastoma SHSY5Y cells in a size-dependent manner.” Int J Nanomedicine 12: 8085–8099.

Livak, K. J. and T. D. Schmittgen (2001). “Analysis of relative gene expression data using realtime quantitative PCR and the 2(-Delta Delta C(T)) Method.” Methods 25(4): 402–408.

Llovera, G., et al. (2014). “Modeling stroke in mice: permanent coagulation of the distal middle cerebral artery.” J Vis Exp(89): e51729.

López-Valdés, H. E., et al. (2014). “Memantine enhances recovery from stroke.” Stroke 45(7): 2093–2100.

Opatz, J., et al. (2009). “SDF-1 stimulates neurite growth on inhibitory CNS myelin.” Mol Cell Neurosci 40(2): 293–300.

Pan, D. S., et al. (2016). “Elevation of serum CXC chemokine ligand-12 levels predicts poor outcome after aneurysmal subarachnoid hemorrhage.” J Neurol Sci 362: 53–58.

Protti, G. G., et al. (2013). “Interleukin-10 may protect against progressing injury during the acute phase of ischemic stroke.” Arq Neuropsiquiatr 71(11): 846–851.

Shyu, W. C., et al. (2008). “Stromal cell-derived factor-1 alpha promotes neuroprotection, angiogenesis, and mobilization/homing of bone marrow-derived cells in stroke rats.” J Pharmacol Exp Ther 324(2): 834–849.

Simkó, M. and M. O. Mattsson (2014). “Interactions between nanosized materials and the brain.” Curr Med Chem 21(37): 4200–4214.

Strle, K., et al. (2001). “Interleukin-10 in the brain.” Crit Rev Immunol 21(5): 427–449.

Szymańska, E. and K. Winnicka (2015). “Stability of chitosan-a challenge for pharmaceutical and biomedical applications.” Mar Drugs 13(4): 1819–1846.

Teleanu, D. M., et al. (2018). “Impact of Nanoparticles on Brain Health: An Up to Date Overview.” J Clin Med 7(12).

Wu, K. J., et al. (2017). “A Novel CXCR4 Antagonist CX549 Induces Neuroprotection in Stroke Brain.” Cell Transplant 26(4): 571–583.

Xu, F., et al. (2013). “Silver nanoparticles (AgNPs) cause degeneration of cytoskeleton and disrupt synaptic machinery of cultured cortical neurons.” Mol Brain 6: 29.

Yin, T., et al. (2013). “SDF-1α in glycan nanoparticles exhibits full activity and reduces pulmonary hypertension in rats.” Biomacromolecules 14(11): 4009–4020.

Yoo, J., et al. (2012). “Effects of stromal cell-derived factor 1α delivered at different phases of transient focal ischemia in rats.” Neuroscience 209: 171–186.

Yuan, Z. Y., et al. (2015). “Brain Localization and Neurotoxicity Evaluation of Polysorbate 80-Modified Chitosan Nanoparticles in Rats.” PLoS One 10(8): e0134722.

Zaman, P., et al. (2016). “Incorporation of heparin-binding proteins into preformed dextran sulfate-chitosan nanoparticles.” Int J Nanomedicine 11: 6149–6159.

Zamproni, L. N., et al. (2017). “Injection of SDF-1 loaded nanoparticles following traumatic brain injury stimulates neural stem cell recruitment.” Int J Pharm 519(1-2): 323–331.

